# Sensitive clustering of protein sequences at tree-of-life scale using DIAMOND DeepClust

**DOI:** 10.1101/2023.01.24.525373

**Authors:** Benjamin Buchfink, Haim Ashkenazy, Klaus Reuter, John A. Kennedy, Hajk-Georg Drost

## Abstract

The biosphere genomics era is transforming life science research, but existing methods struggle to efficiently reduce the vast dimensionality of the protein universe. We present DIAMOND DeepClust, an ultra-fast cascaded clustering method optimized to cluster the 19 billion protein sequences currently defining the protein biosphere. As a result, we detect 1.7 billion clusters of which 32% hold more than one sequence. This means that 544 million clusters represent 94% of all known proteins, illustrating that clustering across the tree of life can significantly accelerate comparative studies in the Earth BioGenome era.

## Main

As the global biosphere is increasingly sequenced and annotated^1,2,3,4,5^, an unprecedented quality of evolutionary insights can now be harnessed to transform the life sciences, where discovery is often driven by the dimensionality reduction of massive experimental data into distinct categories (clusters) that capture common features for inference and predictive tasks. In practice, one such application is the grouping of proteins into related sequence classes that enabled recently celebrated breakthroughs such as protein structure prediction^6,7^, comparative biosphere genomics^8,9^, and classification within metagenomic samples^10,11,12^. These studies are early adopters of leveraging evolutionary information at scale for groundbreaking molecular and functional applications and provide first examples of organizing the entire protein universe for downstream predictive tasks.

Recently, we introduced DIAMOND v2 to meet the user demands for scaling protein search to the tree of life^8^. With DIAMOND v2, we pledged to support the ongoing efforts of the Earth BioGenome project which aims to capture and assemble the genomes of more than 1.8 million eukaryotic species within this decade^3^. In this community quest to compare query sequences against the entire tree of life when millions of species are available, we identified the ability to cluster this vast protein sequence diversity space as a key factor currently limiting the association of sequences across large sets of divergent species.

Here, we perform a comprehensive experimental study to demonstrate that deep-clustering the protein universe of the assembled biosphere which currently consists of ∼19 billion sequences is already possible today. In the Earth BioGenome era, the ability to reduce the sequence space can significantly accelerate protein comparisons when dealing with millions of species and tens of billions of sequences. For this purpose, we estimated that the Earth BioGenome consortium will generate ∼27 billion protein sequences when averaging ∼15,000 genes per species times ∼1.8 million successfully assembled species. Current protein clustering approaches implemented in the standard tools CD-HIT^13^, UClust^14^, and Linclust^15^ are limited when aiming to cluster billions of proteins with such broad sequence diversity in reasonable time and with sufficient clustering sensitivity at lower identity-boundaries. To overcome this limitation and provide a future-proof software solution, we implemented DIAMOND DeepClust, a cascaded clustering method leveraging sensitive protein alignments generated with DIAMOND v2^8^ for incremental clustering and near-complete discovery of distant clusterable homologs at tree-of-life scale (Fig. 1)(Supplementary fig. 1-7). Using DIAMOND DeepClust, we reach this clustering milestone to sensitively cluster 19 billion sequences in 18 days on 27 high performance computing (HPC) nodes (using 250,000 CPU hours in total). This achievement to simultaneously balance speed and sensitivity rather than having to choose between them allows us to substitute Linclust, UClust, and CD-HIT and meet the user demands of the Earth BioGenome project, where clustering sensitivity across large evolutionary distances is paramount. We further optimized our clustering procedure to be memory efficient for laptop users, but also scale linearly in a High Performance Computing (HPC) and Computing Cloud infrastructure to enable sensitive deep-clustering for a vast portfolio of applications (Methods). Finally, we designed an incremental procedure that allows users to add new sequences to a large collection of existing clusters so that the sequencing and assembly community can swiftly add incoming sequences to our biosphere cluster database without the need to re-cluster the entire dataset (Methods).

**Fig. 1.**
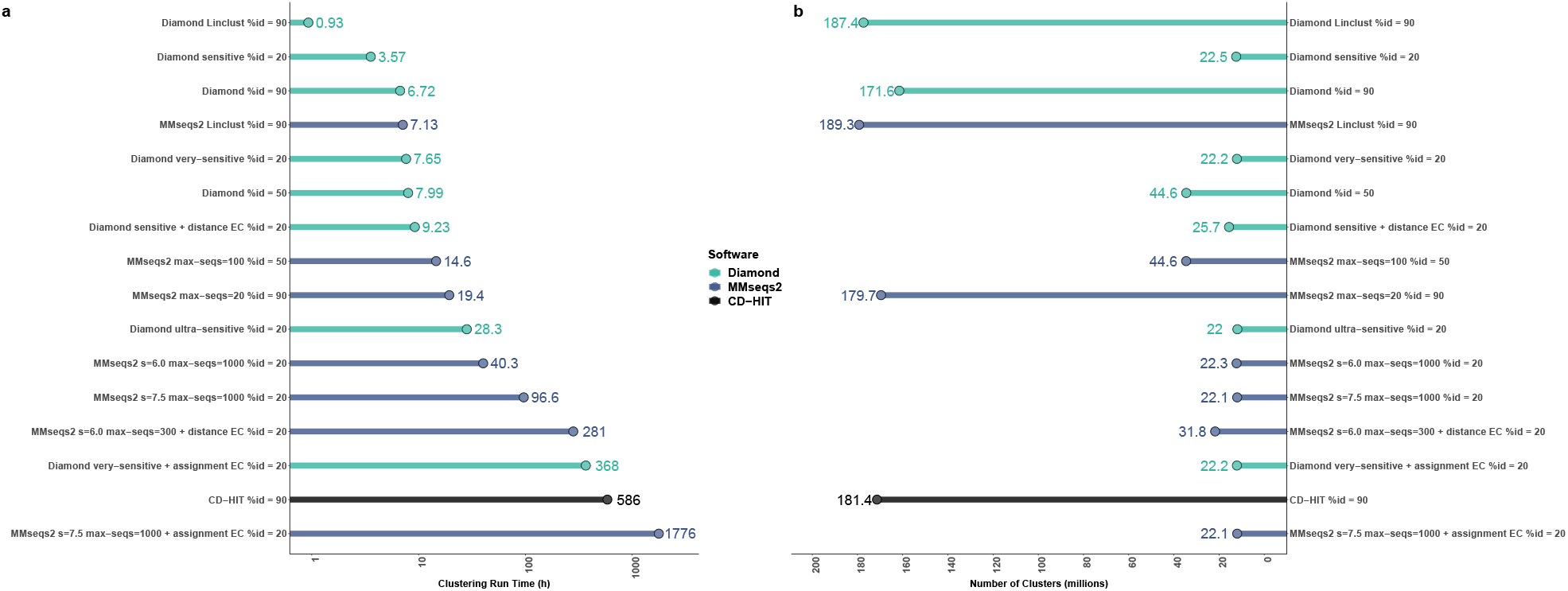
Benchmark of the clustering performance of DIAMOND DeepClust, MMseqs2 and CD-HIT using various sensitivity modes and identity thresholds. Computational benchmarks are shown for clustering the NCBI non-redundant (NR) database currently storing ∼446 million protein sequences using different clustering criteria. **a**, Clustering run times for clustering the NR database on a 64-core server are shown in hours **b**, The resulting cluster counts of compressing the NCBI NR database according to the respective clustering criteria.

As a result of clustering ∼19 billion sequences with 30% sequence identity and 90% coverage thresholds across the tree of life, we determined ∼1.70 billion clusters with 32% of clusters yielding more than one element and 68% denoting singletons (only one unique sequence within each singleton cluster). While this majority of singletons suggests the presence of a large pool of putatively novel proteins (orphan polypeptides) within the protein universe (Experimental Study)(Supplementary fig. 11), these ∼1.16 billion unique sequences comprise only ∼6% of the full set of 19 billion sequences. The fact that 544 million clusters can capture ∼94% of all known proteins illustrates the potential of deep-clustering the protein universe to accelerate protein search across the tree of life.

To put these ∼19 billion sequences of our experimental study into perspective in regard to order of magnitude, we projected that clustering the ∼27 billion eukaryotic protein sequences of the ∼1.8 million Earth BioGenome species with DIAMOND DeepClust would yield ∼2.82 billion clusters and would be feasible today on existing HPC systems (Methods). When assuming similar proportions between singletons (6% of 27 billion sequences) and non-singletons (94% of 27 billion sequences) in this eukaryotic dataset (Supplementary fig. 8-9), we anticipate that the Earth BioGenome project will discover ∼1.6 billion unique (singleton) clusters which can be investigated for their putative molecular function. The overall findings of our experimental study suggest that the protein diversity presently available across all major lineages of life can be reduced by a factor of 10 using sensitive deep-clustering and can further be compressed by a factor of 35 when disregarding singletons, or even by a factor of 60 when removing clusters of size below three. This means that 92% of the protein universe (17.8 billion sequences) can be compressed into 335 million representative sequences for downstream analyses (Supplementary fig. 10). These compression levels can additionally be improved when employing more liberal clustering criteria such as 70% coverage and no identity threshold (compared to our conservative 90% coverage and 30% identity setting). A ProtT5^16^-guided analysis of the protein sequence space mapped by our clustering suggests that it is largely composed of sequences that are mostly uncharacterized by curation efforts such as CATH^17^ or Pfam^18^ (Fig. 2).

**Fig. 2.**
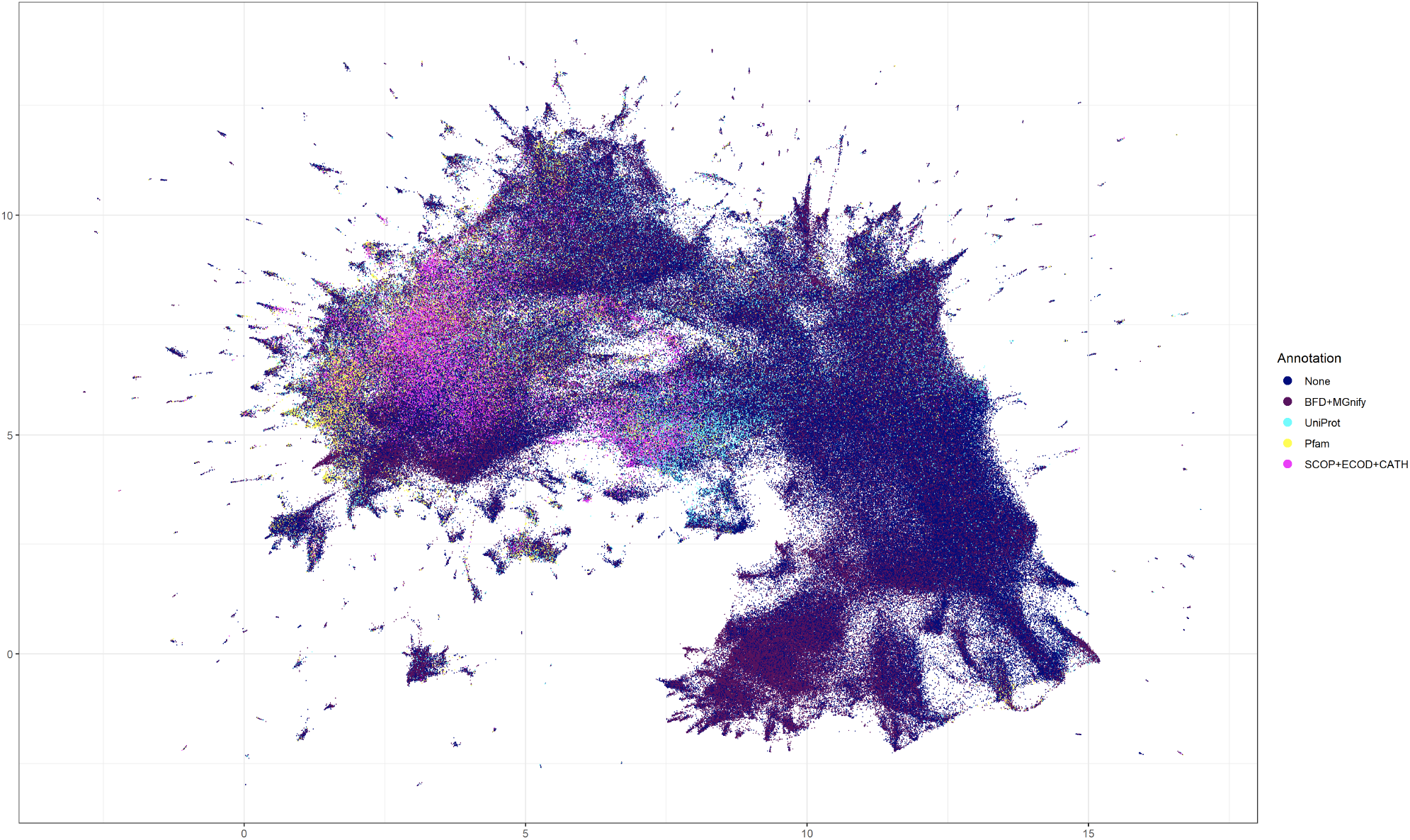
Projection of the protein universe clustered by DIAMOND DeepClust onto its protein language embedding space. UMAP projection of cluster representatives from clusters of size ≥ 5 after transformation into the ProtT5^16^ embedding space. Each dot corresponds to a representative-sequence-embedding labeled by whether the sequence can be annotated with knowledge derived from SCOP^19^+ECOD^20^+CATH^17^ or Pfam^18^, or when no annotation was found whether the respective representative sequence has a homolog in UniProt, or a homolog in the BFD^6^+MGnify^21^ databases. The result illustrates that the protein sequence space is dominated by unexplored protein sequences not sufficiently characterized by standard databases.

Notably, the empowerment enabled by capturing protein diversity across major kingdoms of life, for example, was also demonstrated by recent breakthroughs in protein structure prediction. The predictive power of AlphaFold2 was largely derived from the use of the Big Fantastic Database^6^, a public collection of diverse protein sequences containing 345 million clusters and 61 million clusters with at least three members. While currently holding the status of the largest collection of clustered protein sequences, the result of our experimental study yielding 335 million clusters with at least three members represents a 5.5-fold increase in sequence diversity compared to the Big Fantastic Database which can now be directly incorporated into protein structure prediction research. We therefore provide our 1.7 billion clusters dataset as a free and publicly accessible resource (Data availability).

Algorithmically, DIAMOND DeepClust is based on a cascaded clustering strategy engineered to gradually reduce the complexity of large datasets and to maximally exploit evolutionary conserved information (Methods). Previous methodologies to cluster protein sequences such as CD-HIT^13^ and UClust^14^ are more than ten years old, were not designed to scale to millions of species, and perform poorly when attempting to cluster large datasets deeper than 90% sequence identity. Although MMseqs2/Linclust^15^ presented a considerable advancement over CD-HIT and UClust, it still suffers from comparatively low performance when clustering at high alignment sensitivity, thereby introducing an analytics bottleneck when attempting to scale to >27 billion estimated Earth BioGenome sequences covering the full breadth of biospheric protein space.

To formally benchmark DIAMOND DeepClust against CD-HIT and MMseqs2/Linclust, we clustered the NCBI non-redundant (NR) database containing ∼446M sequences at sequence identity thresholds of 90%, 50% and 20% (Fig. 1). DIAMOND DeepClust solved this problem for deep clustering in 3.6h (sensitive mode) and 7.7h (very-sensitive mode) on a single server equipped with 64 cores compared to 1.7 days and 4 days using MMseqs2, running 11-fold and 13-fold faster respectively. In addition, DIAMOND DeepClust exhibited higher clustering quality as measured by the sensitivity of clusterable homologs found, the completeness of representation by the representative sequences, and the optimality of cluster assignment (Supplementary fig. 2-7). In particular, in the most sensitive run MMseqs2 still did not discover clusterable homologs for 9.8% of the representatives compared to 2.5% for DIAMOND ultra-sensitive (Supplementary fig. 2). Further, for MMseqs2 and DIAMOND, 4.2% of the sequences were not within clustering distance to any representative (Supplementary fig. 3), constituting information that is potentially lost during clustering, while the MMseqs2 workflow to correct such errors increased the runtime to >2 weeks compared to 9.8h using DIAMOND DeepClust (runs labeled as distance error correction), simultaneously degrading the quality and inflating the size of the clustering (cluster count). Lastly, MMseqs2 misassigned 31% of sequences (vs 23% for DIAMOND) to a representative that is not the closest to the sequence (Supplementary fig. 7), while the computation to correct such errors ran for 2.4 months compared to 16 days using DIAMOND DeepClust (runs labeled as assignment error correction). DIAMOND DeepClust ran 82-fold faster than CD-HIT, 3-fold to 8-fold faster than Linclust for clustering at 90% identity and 2-fold faster for clustering at 50% identity.

In conclusion, we designed DIAMOND DeepClust to further optimize the computational steps towards protein alignments against millions of species through dimensionality reduction (clustering) and inspire a new type of research that embraces the biodiversity of life for molecular research and subsequent prediction efforts.

## Acknowledgements

The authors would like to thank Detlef Weigel for his generous sponsorship, discussions, and overall support of this work. We furthermore thank the members of the Drost and Weigel labs for extensive discussions during lab meetings and the Max Planck Computing and Data Facility for providing computational resources. We would also like to thank Jasmin Katz for dedicating her Bachelor Thesis to protein clustering and for valuable discussions in the early phases of this project. Finally, we would like to express our deep gratitude to Alexandru Tomescu, Vikram Alva, Artur Gynter, Fernando Dias, and Andreas Grigorjew, Alexandra Dallaire, Kyanna Ouyang, and Lukas Maischak for trialing early versions of DeepClust in the context of their work and for providing valuable feedback and raise constructive discussions. We thank the BMBF-funded de.NBI Cloud within the German Network for Bioinformatics Infrastructure (de.NBI) (031A532B, 031A533A, 031A533B, 031A534A, 031A535A, 031A537A, 031A537B, 031A537C, 031A537D, 031A538A) for computational support. This work was supported by the Max Planck Society.

## Methods

### Algorithmic overview of DIAMOND DeepClust

#### Representative-based clustering

Following an analogous strategy as the gold standard approaches implemented in CD-HIT^22^ and UCLUST^14^, we define a clustering of an input dataset of protein sequences as a subset of representative sequences such that any input sequence lies within a user-defined distance threshold of at least one representative sequence. As a result, each input sequence will be assigned to one particular representative sequence. This threshold setting (identity, coverage, and e-value) is also referred to as the clustering criterion. For the purpose of this study, in addition to a basic e-value threshold of 0.00001 with respect to the size of the input database, we require that a pairwise local alignment between sequences satisfy a specific minimum sequence identity and length coverage of the co-clustered (non-representative) sequences. Analogous to MMseqs2, we compute the approximate sequence identity instead of the BLAST-like identity defined as the fraction of match columns in the pairwise alignment, as this allows us to save time for backtracing of alignments (Supplementary Information). The approximate identity is derived from the alignment score and the lengths of the aligned ranges in the sequences as a linear regression and can be considered a better measure of evolutionary distance than the actual sequence identity^23^. Starting from a set of alignments that meet the user-specified or default clustering criterion, we compute a set of representative sequences by first encoding the alignments as a directed graph G where nodes represent individual protein sequences and edges denote pairwise local alignments between them, whereby a directed edge from sequence A to sequence B indicates that A can represent B according to the clustering criterion. In a second step, we apply the greedy vertex cover algorithm on the alignment-graph G to determine a near-minimal covering set of graph vertices. The algorithm repeatedly selects the vertex with the highest node outdegree and removes it from the graph along with its out-neighbors to form a new cluster until the graph is completely clustered. In the final round of cascaded clustering, we also permit recursive merges of clusters to prevent clusterable pairs to remain in the final clustering. We also implemented simple length-sorted clustering but observed better clustering quality for the greedy vertex cover approach (data not shown).

#### Cascaded clustering

As exhaustive all-vs-all alignment of protein datasets consisting of hundreds of millions of sequences is prohibitively expensive, we approach this issue by adopting cascaded clustering^24^ to gradually construct larger sequence clusters in several rounds of comparison with increasing alignment sensitivity (iterating between modes: --fast, *default*, --sensitive, --very-sensitive, --ultra-sensitive). In the first round, we subsample the seed space using minimizers^25^ with a window size of 12 which we empirically found to provide a good balance between speed and sensitivity, and attempt to achieve linear computational scaling of comparisons by considering only seed hits against the longest sequence for identical seeds rather than trialing all possible combinations^15^. This heuristic is sufficient to find meaningful representatives as the seeds at this stage are selected to be highly specific and the longest sequence is a priori the most likely to maximize recruitment of member sequences due to the unidirectional length coverage criterion. We compute representative sequences from the resulting alignments by greedy vertex cover^24^, which are then passed on to the next round of cascaded clustering and subjected to an all-vs-all DIAMOND v2 blastp search at increased sensitivity. Depending on the desired clustering depth, two to six of these alignment rounds are chained until reaching sufficient sensitivity such that most representatives within clustering distance have been discovered. We optimized self-alignment of the representative databases by taking advantage of the symmetry of queries and targets in the seeding stage, avoiding the evaluation of redundant seed hits, and thus doubling the performance of this computation. This is accordingly taken into account when the database is processed in blocks by eliminating redundant block combinations.

#### Distance error correction

Distance errors in the clustering are introduced by sequences that do not fall within the clustering distance of their assigned representative and arise due to the recursive merging of clusters in the cascaded clustering workflow based on alignments of only the representative sequences. These errors do not necessarily present an error in the biological sense, since biological properties of the sequences such as ancestry, structure and function can be conserved despite the fact that the local alignment does not satisfy a certain threshold requirement. Nevertheless, we implemented an additional workflow to optionally correct such errors. We first align all sequences against their assigned representatives to find the sequences failing the clustering criterion.

After the identification of all putatively mis-clustered sequences, we re-align the set of these sequences against the database of all representative sequences using DIAMOND v2 in iterated blastp search mode with increasing sensitivity. If an alignment against a representative sequence is detected, the sequence is reassigned to the cluster of that representative. If multiple representatives satisfy the clustering criterion, the e-value of the local alignment determines the assignment. We collect all sequences that fail to align against any representative, remove them from the clustering and re-cluster this dataset with the cascaded clustering workflow. The resulting sub-clustering is again subjected to the distance error correction workflow, an iterative procedure that continues until convergence to a clustering with no distance errors.

#### Assignment error correction

For any given clustering a sequence may lie in clustering distance of multiple representative sequences. Cascaded clustering or incremental clustering as performed by tools like CD-HIT or UClust hold no guarantee of assigning a sequence to the cluster of the closest representative, measured by a metric such as the e-value of the local alignment. Although this property of assignment error has no impact when users wish to work only with the set of representative sequences in the context of dimensionality reduction, it becomes a relevant drawback when attempting to use all cluster members for downstream analyses such as multiple sequence alignments or gene family characterization. To this end, we have implemented a reassignment workflow that will search all non-representative sequences against the representative database and assign each sequence to the closest representative as measured by the e-value (if the e-value is 0 for different representatives the bitscore determines the representative assignment), while maintaining the clustering criterion.

#### Many-core parallelization of clustering tasks

Analogous to our computational scaling efforts introduced in DIAMOND v2, we elevated our clustering capabilities to run massively parallel on High-Performance-Computing (HPC) and Cloud Computing infrastructures. To accommodate servers with 128 or more compute cores, we have refactored our multi-threading code to fix existing load imbalances during the alignment workflow and allow the software to scale smoothly to 256 threads or more. Optimal scaling of the seed extension stage in DIAMOND v2 is impeded by query proteins that attract a disproportionate number of target hits or incur an unusual cost of Smith Waterman extensions based on the length of their respective sequence, due to the use of static load balancing that is only able to distribute different query sequences among threads. We addressed this issue in DIAMOND DeepClust by implementing a fine granular task-based parallelism in which individual threads that are processing expensive queries can make use of a work-stealing task scheduler to redistribute extension tasks among the thread pool.

#### Gapped alignment computation

We produced a novel vectorized Smith Waterman implementation based on a modified SWIPE^26^ approach that was originally developed for the first DIAMOND version^27^, but dropped out of its code base soon after the initial release. While SWIPE vectorized the Smith Waterman algorithm by computing alignments of the same query against multiple targets, we generalize this approach to computing alignments of multiple independent query/target pairs. This is accomplished by using score profiles for the queries that store alignment scores along the sequence for each of the amino acid residues. For computing one column of the DP matrix, we maintain pointers into these profiles for each SWIPE channel and apply an AVX2-optimized matrix transposition to interleave the query/target scores for each DP cell into the same register, then compute the cell updates according to the standard SWIPE logic. Contrary to the original SWIPE design, this approach permits the computation of banded and anchored alignments, which in turn also enable optimizations such as the cheap determination of alignment start and end coordinates as well as X-drop termination. Compared to our implementation of the original SWIPE algorithm, we have measured ∼20% computational overhead for this approach on the Intel Ice Lake architecture.

#### Ungapped alignment heuristics

When clustering highly similar sequences at >90% identity, most ungapped segment pairs that make up their alignments can already be found during the seeding stage. We exploit this by clustering sequences without full Smith Waterman extension if one of the ungapped alignments emerging from the seed hits already satisfies the clustering criterion. Conversely, we exclude a target from gapped extension if the sum of identities or sequence coverage of these ungapped segment pairs fails a relaxed clustering criterion based on empirically derived thresholds. The first heuristic for accepting alignments without full extension was also used for the clustering runs at 50% sequence identity. Together, this ungapped alignment strategy allows to reduce the computational burden when dealing with highly similar sequences without the loss of clustering sensitivity.

#### Memory optimization

Next to speed, sensitivity and user-friendliness, feedback from DIAMOND users identified memory efficiency of the search procedure as one the main advantages compared to alternative aligners. This memory optimization feature of DIAMOND allows users to perform large scale searches on their laptops and scale their parallelization efforts seamlessly into an HPC or cloud infrastructure through its distributed memory and parallelization library. Our aim for DIAMOND DeepClust was therefore to continue this memory efficiency streak by designing a memory efficient cascaded clustering to unlock the clustering of large input datasets on a laptop or massively parallel virtual machines when aiming to scale distributed computing in the cloud. In detail, the double indexing approach with runtime-generated and partitioned indexes allows the aligner to operate memory-efficiently without the need to store large index data structures on disk or maintain them in memory. For clustering large datasets with limited memory, DIAMOND will automatically use an incremental procedure on a length-sorted and partitioned database as described under Experimental Study/Clustering, which effectively limits both the use of temporary storage space and main memory to a user-defined maximum (command line option -M). For clustering on HPC systems, the design of the cascaded clustering algorithm as chained rounds of all-vs-all alignments at increasing sensitivity also allows decomposition of this computation into arbitrarily many work packages that can be processed independently on a distributed infrastructure. This process can be automated using the multiprocessing feature introduced in DIAMOND v2^8^.

#### Cluster extension

Sensitively clustering billions of sequences across the tree of life is a computationally heavy task. However, this procedure has to be performed only once and the resulting cluster database can be made publicly available (Data availability). To accommodate the future growth of sequenced species within this decade, we designed a cluster extension workflow to allow users to add new query sequences to existing clusters to extend the initial cluster database without the need of re-clustering all sequences together. In particular, new sequences can be searched against the existing representative set using DIAMOND in iterative search mode (option --iterate). This mode identifies all sequences that can be assigned to an existing cluster. The remaining unaligned sequences can then be clustered independently, and the resulting representatives can be added to the existing clustering.

## Benchmarks

**Supplementary fig. 1.**
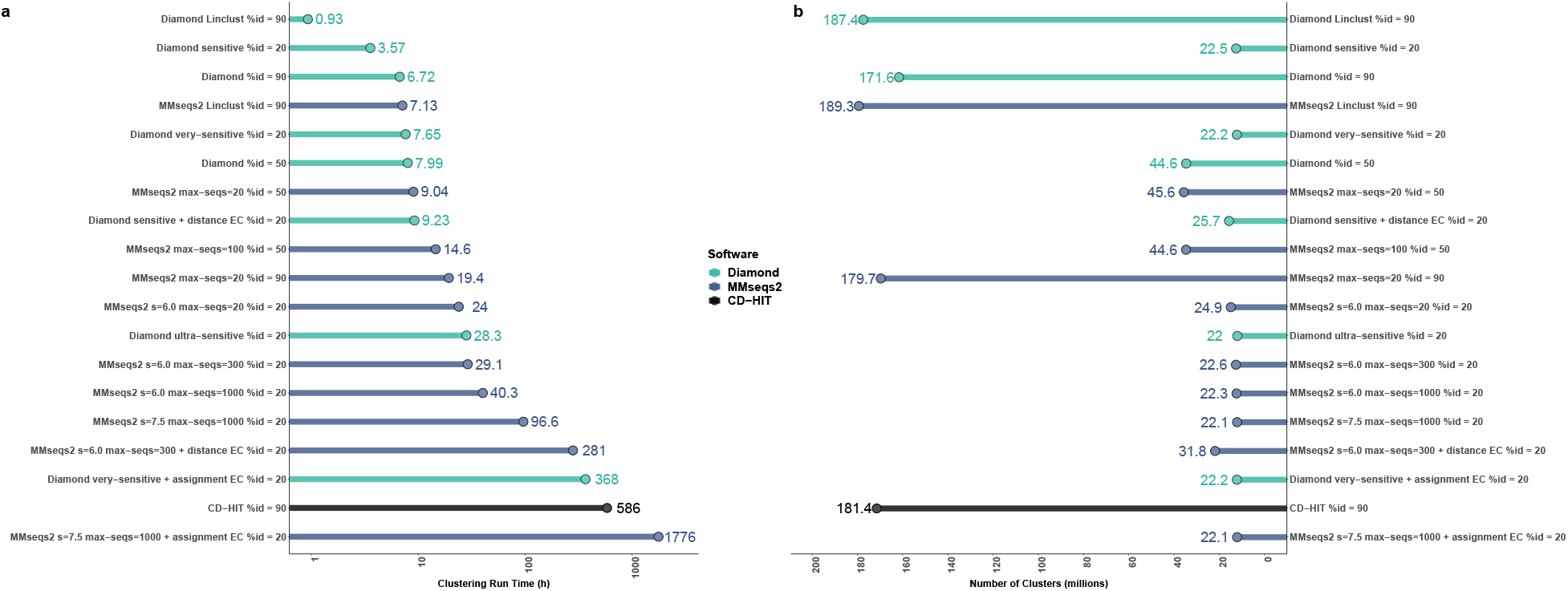
Benchmark of the clustering performance of DIAMOND DeepClust, MMseqs2 and CD-HIT using a broader variety of max-seqs parameters. To cover a broader range of parameter settings, we benchmarked various alternative settings of the MMseqs2 max-seqs parameter against DIAMOND DeepClust and CD-HIT. **a**, Run times for clustering the NCBI NR database on a 64-core server are shown in hours **b**, An illustration of the cluster counts resulting from clustering the NCBI NR database with the respective clustering criteria.

### Design

Our clustering benchmark is based on the NCBI NR database downloaded in November 2022, containing 513,991,389 sequences and 200,929,118,620 total residues. We hard-masked this database using tantan^28^ with default settings and removed all sequences that were masked over >10% of their range, resulting in a reduced database of 445,610,930 sequences. This choice is motivated by the fact that stringent filtering of false positives is important for deep clustering, which is normally handled by DIAMOND and MMseqs2 by applying soft-masking and composition-based score correction. For the purpose of our benchmark, we did not want to rely on the tools’ internal masking and score correction procedures however, as this would impair the comparability of results. We therefore disabled these features for our benchmarking runs and instead relied on the precomputed hard masking. Removing sequences that are masked over a substantial part of their range is necessary for this design since otherwise they would remain as unclusterable singletons throughout the computation due to the coverage criterion, needlessly inflating the runtime. For each of the benchmarked tools, we report the wall clock time for clustering this database and the resulting number of clusters. We conducted separate runs with a clustering criterion of 90% and 50% sequence identity, as well as no restriction of the sequence identity, while setting a coverage cutoff of 80% of the co-clustered sequence for all runs. The deep clustering runs without restriction of the sequence identity were executed three times using the --sensitive, --very-sensitive and --ultra-sensitive modes of DIAMOND, as well as the -s6.0 and - s7.5 sensitivity settings of MMseqs2, which we chose to roughly correspond to the first two DIAMOND modes^8^. The --max-seqs parameter of MMseqs2 needs to be manually set by the user, so we decided to try a set of possible values whereby further increases were limited by the amount of disk space available on the benchmark system (Supplementary fig. 1). We selected the run that was most comparable to the corresponding DIAMOND run based on the sensitivity error metric (Supplementary fig. 2) for creating (Fig. 1). We limited the evaluation of CD-HIT to clustering at 90% identity since deeper clusterings at lower sequence identity levels cannot be computed with this tool in practical time. For DIAMOND and MMseqs2, we conducted additional runs labeled as *distance error correction*, designed to correct errors where sequences do not satisfy the clustering criterion against their assigned representative. These runs correspond to the recluster workflow in DIAMOND and the --cluster-reassign option of MMseqs2. For DIAMOND and MMseqs2, we conducted additional runs labeled as *assignment error correction*, designed to reassign each non-representative sequence to the cluster of the closest representative (as measured by the e-value of the local alignment) that satisfies the clustering criterion. The actual computations were distributed on a compute cluster and the runtimes converted to the equivalent of a single 64-core server. The benchmarks for clustering at 90% identity were run based on an older version of the NR database downloaded in September 2021 containing 425,032,034 protein sequences and 155,806,124,097 total residues. The database was not hardmasked and no sequences were excluded.

### Environment

All runs were conducted on a pair of 64-core dedicated virtual cloud nodes with 1 TB RAM and a 2 TB SSD on the HPC Cloud at the Max Planck Computing and Data Facility in Garching. The hypervisors used were dual Intel IceLake-based (Xeon Platinum 8360Y @ 2.4 GHz) compute nodes with a total of 72 cores, 2 TB of RAM and 10 TB SSD storage in RAID 6 configuration. The HPC Cloud is based on OpenStack and CEPH and offers standard cloud computing “building blocks”, including virtual machines based on common Linux operating systems, software-defined networks, routers, firewalls, and load balancers, as well as integrated block and S3-compatible object storage services. Analogous infrastructures are currently employed by centralized commercial cloud computing providers such as Google Cloud, Amazon Web Services, and Microsoft Azure, thereby allowing native adoption of DIAMOND DeepClust on these systems as well.

**Supplementary fig. 2.**
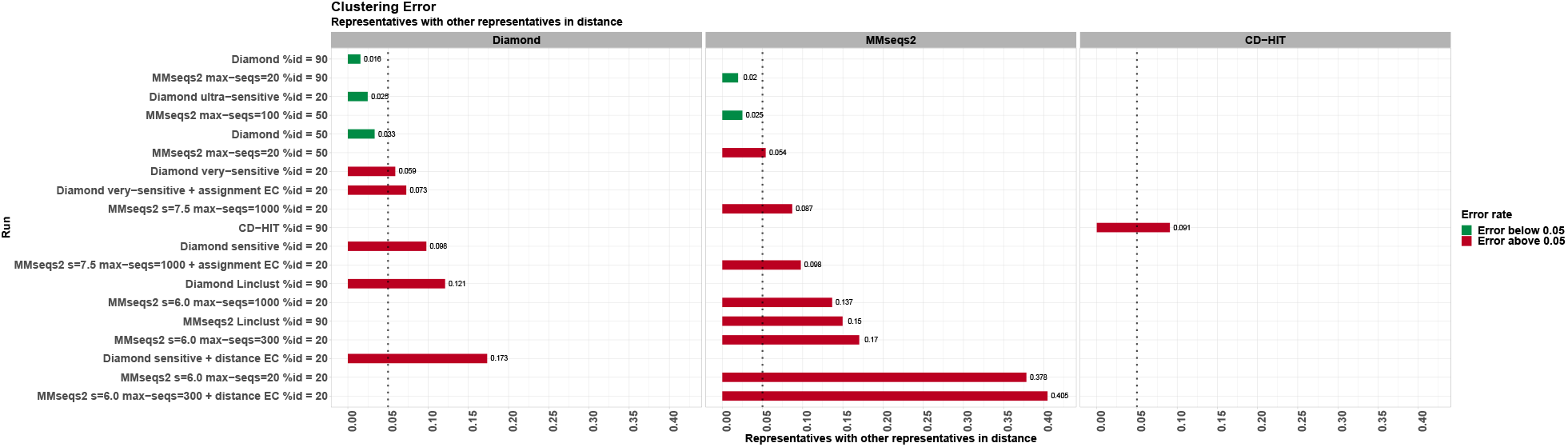
Benchmark of the sensitivity errors with DIAMOND DeepClust and MMseqs2 using various clustering criteria. Shown is the fraction of cluster representative sequences that satisfy the clustering criterion against another representative. Error rates below 0.05 indicate that most representatives are unique in the clustered set and do not correspond to other representatives.

**Supplementary fig. 3.**
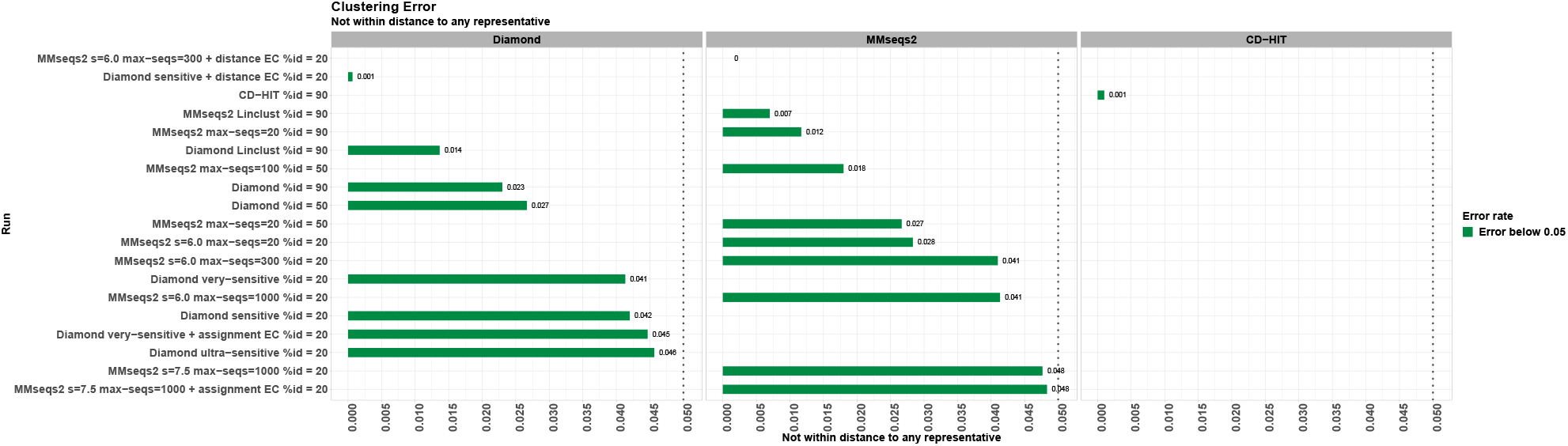
Benchmark of distance errors when clustering with DIAMOND DeepClust and MMseqs2 using various clustering criteria. Shown is the fraction of cluster member sequences that do not satisfy the clustering criterion against any representative sequence.

**Supplementary fig. 4.**
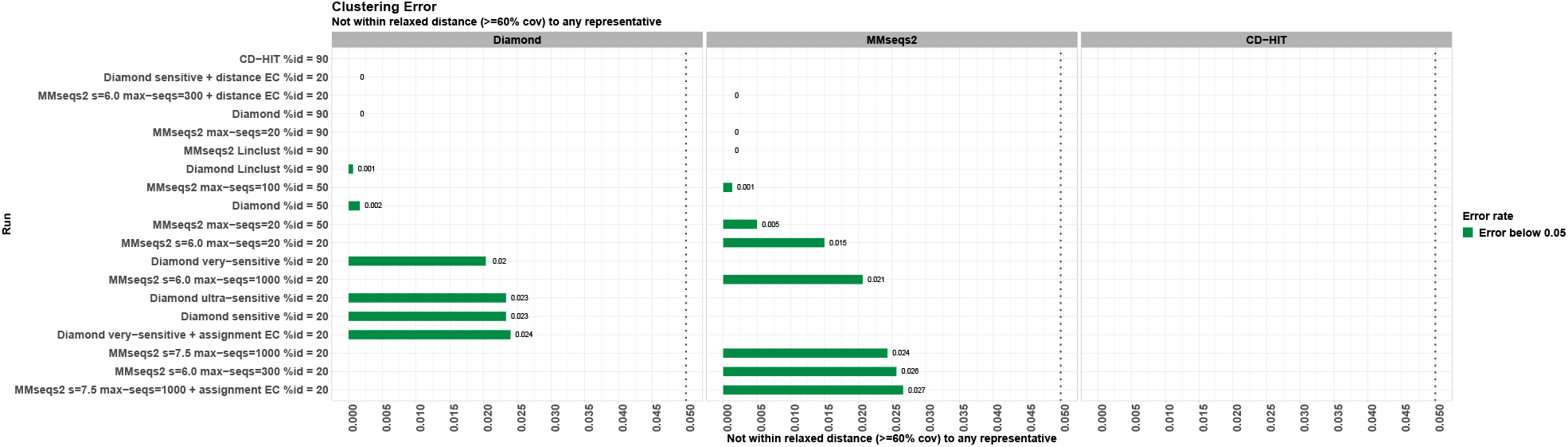
Benchmark of distance errors (relaxed) when clustering with DIAMOND DeepClust and MMseqs2 using various clustering criteria. Shown is the fraction of cluster member sequences that do not satisfy a relaxed clustering criterion against any representative sequence, defined as 60% coverage and a sequence identity threshold lowered by 10 percentage points.

**Supplementary fig. 5.**
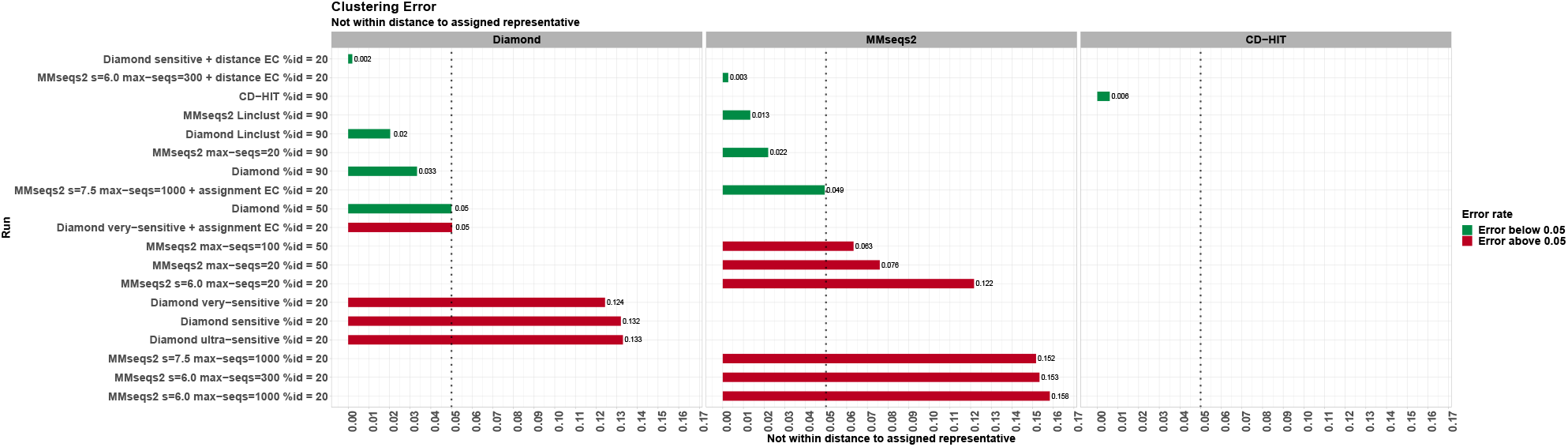
Benchmark of distance errors when clustering with DIAMOND DeepClust and MMseqs2 using various clustering criteria. Shown is the fraction of cluster member sequences that do not satisfy the clustering criterion against their assigned representative sequence.

**Supplementary fig. 6.**
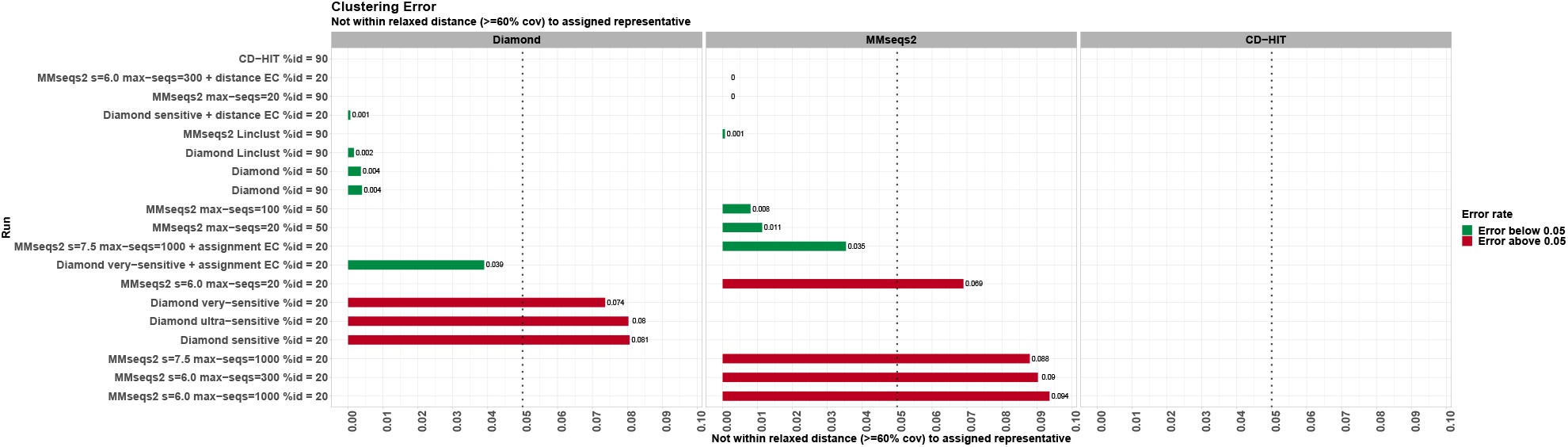
Benchmarking of distance errors (relaxed) when clustering with DIAMOND DeepClust and MMseqs2 using various clustering criteria. Shown is the fraction of cluster member sequences that do not satisfy a relaxed clustering criterion against their assigned representative sequence, defined as 60% coverage and a sequence identity threshold lowered by 10 percentage points.

**Supplementary fig. 7.**
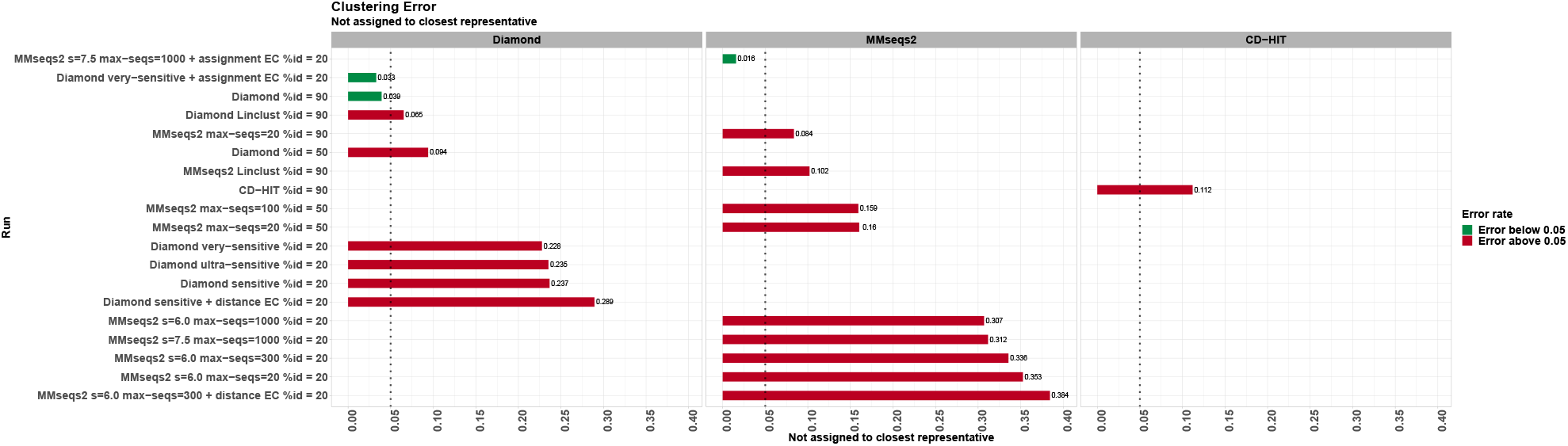
Benchmark of assignment errors when clustering with DIAMOND DeepClust and MMseqs2 using various clustering criteria. Shown is the fraction of cluster member sequences that are not assigned to the closest representative that satisfies the clustering criterion, defined by the e-value of the local alignment. Smaller error rates indicate that most cluster members are indeed sufficiently assigned to their closest representative and do not match other (closer) representatives better than the assigned representative.

### Clustering evaluation

We show four different metrics to measure errors in the computed clusterings. First, we show the fraction of representative sequences that fulfill the clustering criterion against another representative and represent false clusters that could have been merged into one unifying cluster but were missed due to limited alignment sensitivity (Supplementary fig. 2). Second, we show the fraction of cluster member sequences that do not satisfy the clustering criterion against their assigned representative, errors that are caused by the cascaded clustering algorithm (see distance error correction)(Supplementary fig. 5-6). Third, as a relaxed version of the second metric, we show the fraction of cluster member sequences that do not satisfy the clustering criterion against any representative and therefore constitute information that is potentially lost in the representative set (Supplementary fig. 3-4). Fourth, we report the fraction of cluster member sequences that are not assigned to the closest representative that satisfies the clustering criterion, as measured by the e-value of the local pairwise alignment (Supplementary fig. 7). The cascaded clustering algorithm as well as the incremental algorithm used by CD-HIT may both produce such suboptimal assignments by design, causing errors in the clustering that could be undesirable depending on the application (see assignment error correction).

We established the ground truth for these evaluations by computing a full Smith Waterman alignment of the evaluated representative or cluster member sequences against all representative sequences using DIAMOND in --swipe mode which guarantees perfect pairwise alignment sensitivity. Due to the much larger representative set, we used DIAMOND in default mode for the 90% identity runs. On account of the expense of computing the exhaustive Smith Waterman alignments, we evaluated these error metrics on random samples of 3,000 representative and cluster member sequences respectively. We sampled 3,000 clusters for each run from the set of clusters containing at least 5 sequences, and additionally sampled one member sequence out of each of these clusters. Since this evaluation is based on comparing raw alignment output, we added an option to DIAMOND to mimic the alignment score computations of MMseqs2 (Supplementary Information). We computed 95%-confidence intervals for the error metrics based on the procedure of Clopper and Pearson^29^ (source data for Supplementary fig. 2-7).

**Supplementary Fig. 8.**
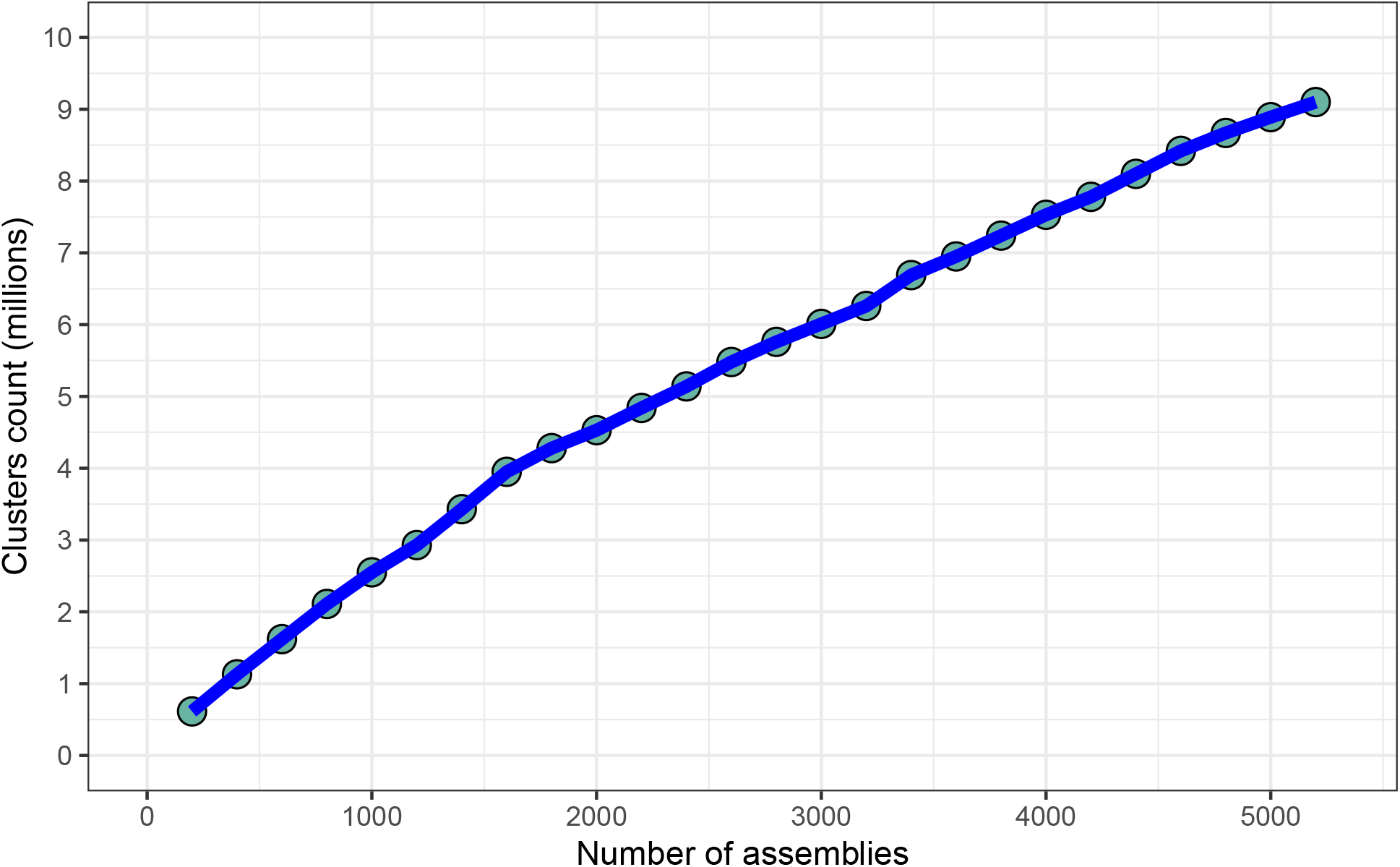
Eukaryotic protein clusters generated with DIAMOND DeepClust. Shown is the number of clusters resulting from clustering all 77.3 million protein sequences derived from up to 5,155 eukaryotic assemblies. The monotonically increasing graph illustrates that the eukaryotic sequence diversity space is not fully saturated yet, suggesting that the efforts of the Earth BioGenome Project have the potential to add a sufficient proportion of eukaryotic protein diversity to the existing protein universe. Number of clusters (millions)

**Supplementary Fig. 9.**
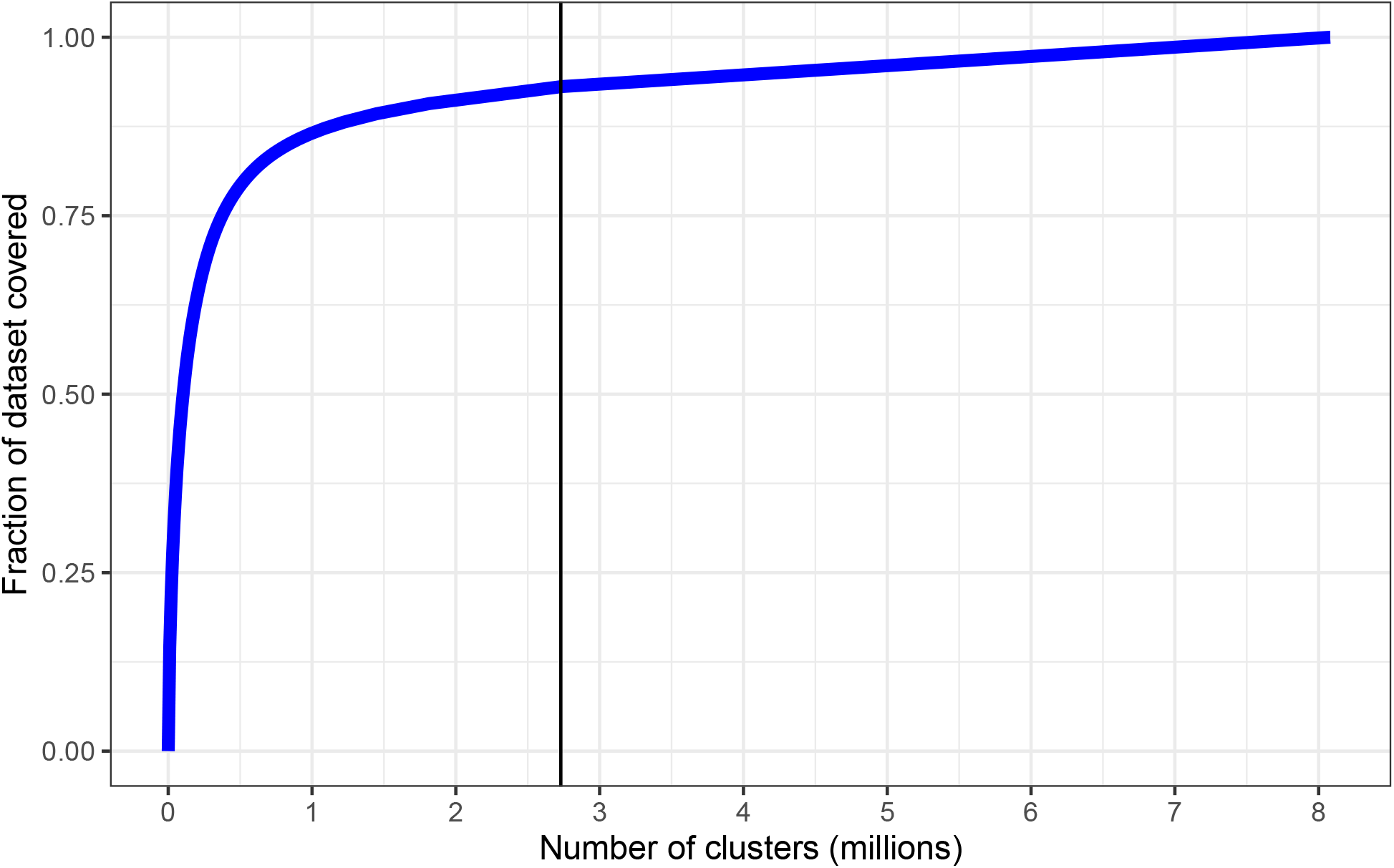
Cluster representation for eukaryotic proteins from clusters generated with DIAMOND DeepClust. Shown is the number of clusters against the fraction of the 77.3 million eukaryotic proteins that are covered by the clustering. The vertical line illustrates that eukaryotic sequences cluster in similar proportions between non-singletons (93%) vs singletons (7%) as the full protein universe (∼19 billion sequences; 94% non-singleton vs 6% singleton) which is currently dominated by protein sequences derived from microbial samples.

### Projection for Earth BioGenome Era

Clustering the protein universe of our earth’s biosphere allows us to quantify and understand the complexities and degrees of divergence when aiming to apply the comparative method across the tree of life and harness insights of relatedness for downstream analyses. To estimate the future applicability of DIAMOND DeepClust, we projected the computational effort of running it on a future dataset that would cover all protein sequences retrieved from the ∼1.8 million eukaryotic species expected to be sequenced by the Earth BioGenome Project. To this end, we downloaded the protein sequences of 5,155 eukaryotic assemblies with annotated genes that were available in GenBank as of July 2022 and randomly partitioned them into groups of 200 assemblies. We clustered the protein sequences of the first group using DIAMOND DeepClust in very-sensitive mode at a 75% coverage cutoff and no identity cutoff and kept adding the sequences of another group to the clustering as described above under Cluster extension, until all assemblies were added. We observed a linear growth of the cluster count in the number of species with no apparent saturation (Supplementary fig. 8). Based on a number of 8,080,544 clusters for this dataset, we could project a linear growth of the cluster count as an upper bound estimate, resulting in 2.82 billion clusters for 1.8 million species. The clustering computation is dominated by all-vs-all alignment in the most sensitive clustering round and can thus be assumed to scale roughly quadratically in the number of clusters. On the basis of this projection assumption and a DIAMOND v2 runtime of 2.07h on a 72-core server for the given computation, we project a computation time of 18.1 million CPU hours for processing the full Earth BioGenome dataset with DIAMOND DeepClust. Such a computation is already feasible on HPC systems hosted by the Max Planck Society today and illustrates that the scalability of DIAMOND DeepClust will enable users to learn from the protein sequences of millions of species once they are available. We note that the actual cluster count and run time will likely be lower due to saturation effects. While the nature of prokaryotic versus eukaryotic gene expression and regulation is fairly different, our clustering of all available eukaryotic proteins shows that after clustering the ratio between singletons (66%) vs non-singletons (34%) is comparable to our microbes dominated dataset of 19 billion sequences in that non-singleton clusters capture 93% of all eukaryotic proteins (Supplementary Figure 9).

**Supplementary fig. 10.**
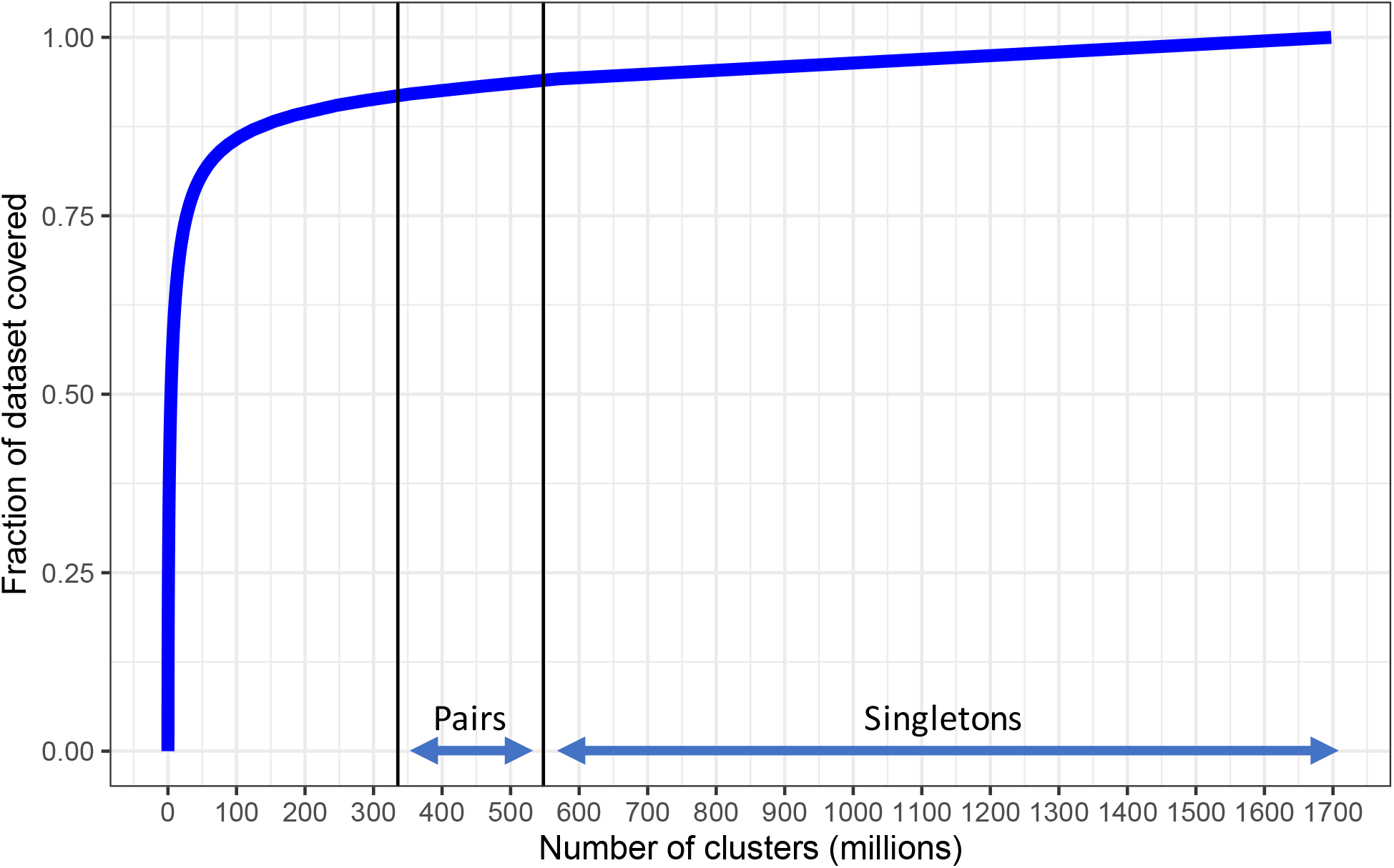
Cluster representation for the experimental study. Shown is the number of clusters generated with DIAMOND DeepClust against the fraction of the 19.4 billion input proteins that are covered by the clustering. The vertical lines indicate the start of clusters of size two and one (from left to right). The result illustrates that ∼335 million representatives can capture 92%(∼ 17.48 billion sequences) of the protein universe that comprises 19 billion sequences.

### Experimental study

Recently, we introduced DIAMOND v2 to unlock the familiar functionality of a BLASTP search for tree-of-life scale applications in the Earth BioGenome era. To mimic protein alignments at this scale, we aligned ∼280 million sequences from the NCBI NR database against ∼39 million sequences of the UniRef50 database which resulted in ∼32 billion pairwise alignments which could be performed in less than 6h on a HPC (compared to several months with BLASTP) while matching the sensitivity of BLASTP^8^. While this speedup allowed us to introduce DIAMOND v2 as biosphere-ready protein aligner, searching protein sequences against millions of species and yielding trillions of pairwise alignments remains a computational challenge. Overcoming this bottleneck requires extensive dimensionality reduction of protein sequence space into sequence clusters to perform pairwise alignments only on the set of representative sequences rather than all sequences.

Using DIAMOND DeepClust, we performed an experimental study to showcase the power of dimensionality reduction through sequence clustering when the Earth BioGenome project will have successfully sequenced and assembled all ∼1.8 million species. For this purpose, we collected ∼22 billion protein sequences across all kingdoms of life (currently mostly comprising of microbes) to match the order of magnitude and sequence diversity space of the estimated ∼27 billion eukaryotic protein sequences planned to be generated as part of the Earth BioGenome project (assuming ∼1.8 million species times an average of ∼15,000 genes per species). We note that compared to our collection of ∼22 billion sequences (∼19.4 billion deduplicated sequences), the Earth BioGenome set will include a much broader sequence diversity space derived from eukaryotes (while our current dataset is enriched in proteins mostly derived from microbial metagenomic samples).

**Supplementary fig. 11.**
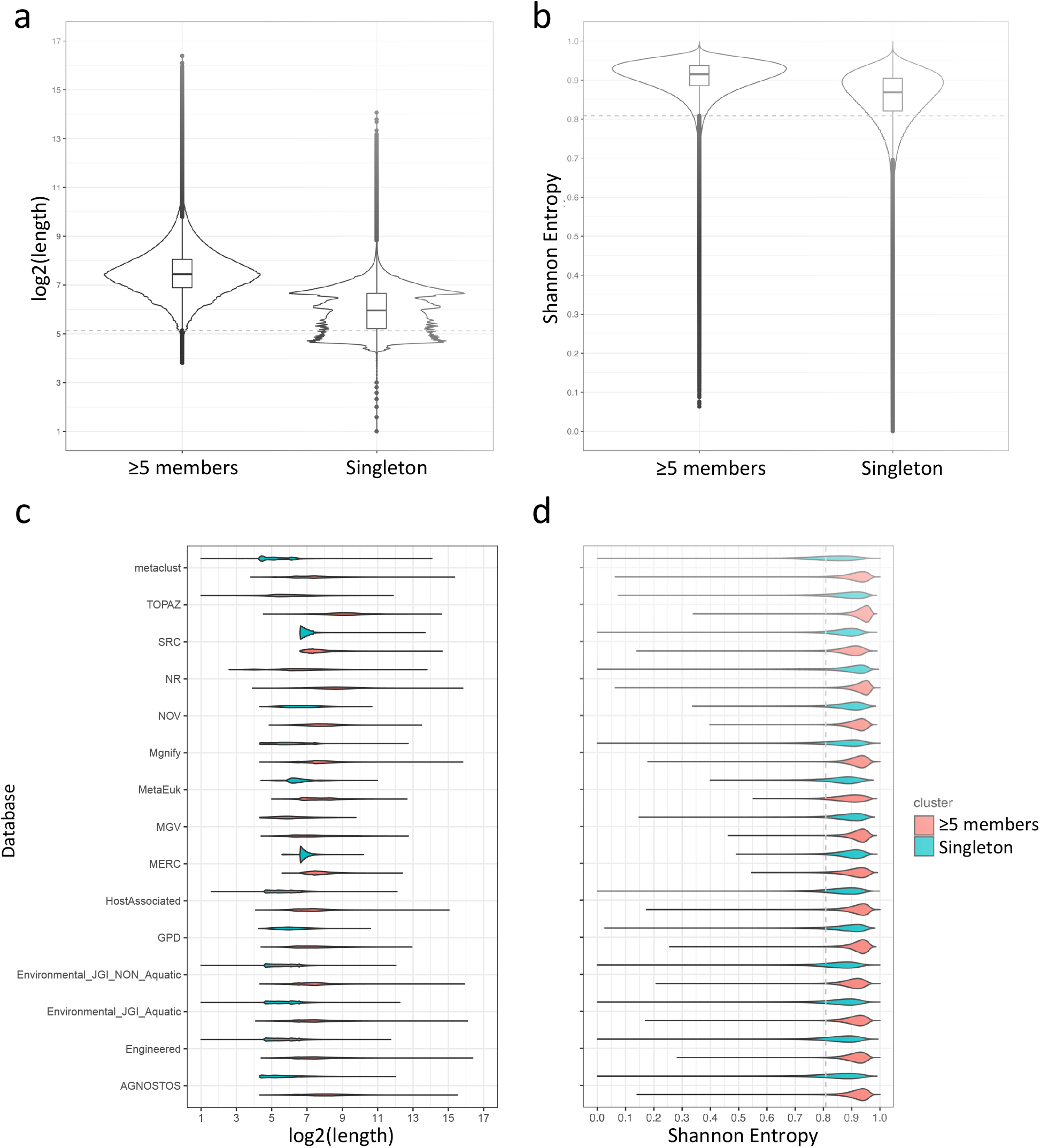
Characterization of the 1.7 billion clusters generated by DIAMOND DeepClust. Illustrated are sequence length and Shannon entropy for cluster representatives. The length and Shannon entropy (sequence randomness) were computed for the representatives of clusters with at least five members compared with the singletons. IQR outlier cutoff was determined based on the distribution computed for clusters with at least five members, denoted as dashed lines. **a**. Sequence length distribution of non-singleton versus singleton clusters, **b**. Shannon entropy distribution of non-singletons versus singleton clusters. **c**. Sequence length distributions shown in **a**, now grouped by the public database the respective sequence originated from. **d**. Shannon entropy distributions shown in **b** now grouped by the public database the respective sequence originated from.

As a result of clustering ∼19 billion deduplicated sequences with 30% sequence identity and 90% coverage thresholds across the tree of life, we determined ∼1.70 billion clusters with 32% of clusters yielding more than one element, 12% of clusters holding more than five elements, and 68% of clusters with only one element (singletons)(Supplementary fig. 10). This striking result shows that while 68% of clusters contain unique sequences, these ∼1.16 billion singletons represent only 6% of the 19 billion sequences defining our current protein universe which begs the question whether these distinct proteins are derived from novel orphan genes or whether they represent assembly and annotation artifacts. To inform ongoing mega-assembly consortia about the proportion of novelty versus assembly artifact we compare the sequence and protein properties of singletons vs non-singletons. First, we filtered out obvious artifacts and poorly annotated sequences by running the repeat masking software tantan^28^ with default settings on a random sample of 1,000,000 singletons vs non-singletons to evaluate the proportion of low-complexity sequences among putatively novel proteins. Interestingly only 10.5% of proteins were masked over >25% of their range by tantan compared to 9.41% for non-singletons. Next, we compare the length distributions of sequences derived from singleton clusters vs non-singleton clusters, since a well-studied feature of novel proteins is their short average length compared to evolutionarily conserved proteins^30^. As a result, we find that singletons are indeed significantly shorter than non-singleton protein sequences with singletons comprising a median length of 62 amino acids and non-singletons 174 amino acids (Supplementary fig. 11). In addition to sequence length, the entropy in the protein sequence is also known to vary between novel and evolutionarily established proteins^30^. Therefore, we calculated the Shannon entropy for all singletons vs non-singletons and observed that singletons indeed show significantly higher entropy values than non-singletons, thereby illustrating the uneven nature of these unique novel proteins (orphan polypeptides) (Supplementary fig. 11). Furthermore, we aligned random samples of 1,000,000 singletons vs 1,000,000 non-singletons (from clusters containing ≥ 5 members) against the ECOD database^20^ using DIAMOND in ultra-sensitive mode and found that 11,601 sequences (∼1.16%) generated alignments with e-value < 0.001, while 180,421 non-singleton sequences (∼18%) generated alignments with the same alignment settings. This fact further establishes that the ∼1.16 billion singletons we report in this study require further attention to assess their biological relevance. Since these singletons show signatures previously assigned to novel orphan proteins they encourage further studies. For example, the clustering run used in creating the AlphaFold2 Big Fantastic Database used only 18% of all clusters and constrained the cluster size to at least three elements, while removing 82% of singleton clusters. If, however, further research would reveal that they are the product of poor mega-assembly efforts, our current representation of the protein universe would turn out to be heavily biased by assembly quality, particularly when derived from metagenomic assemblies. Together, our experimental study reveals an unprecedented quantification of the protein universe and will enable future efforts to test whether more high quality genome assemblies based on long-read technologies will yield smaller numbers of singleton clusters when joining the tree of life or whether this unique diversity is an intrinsic feature of life itself.

**Supplementary fig. 12.**
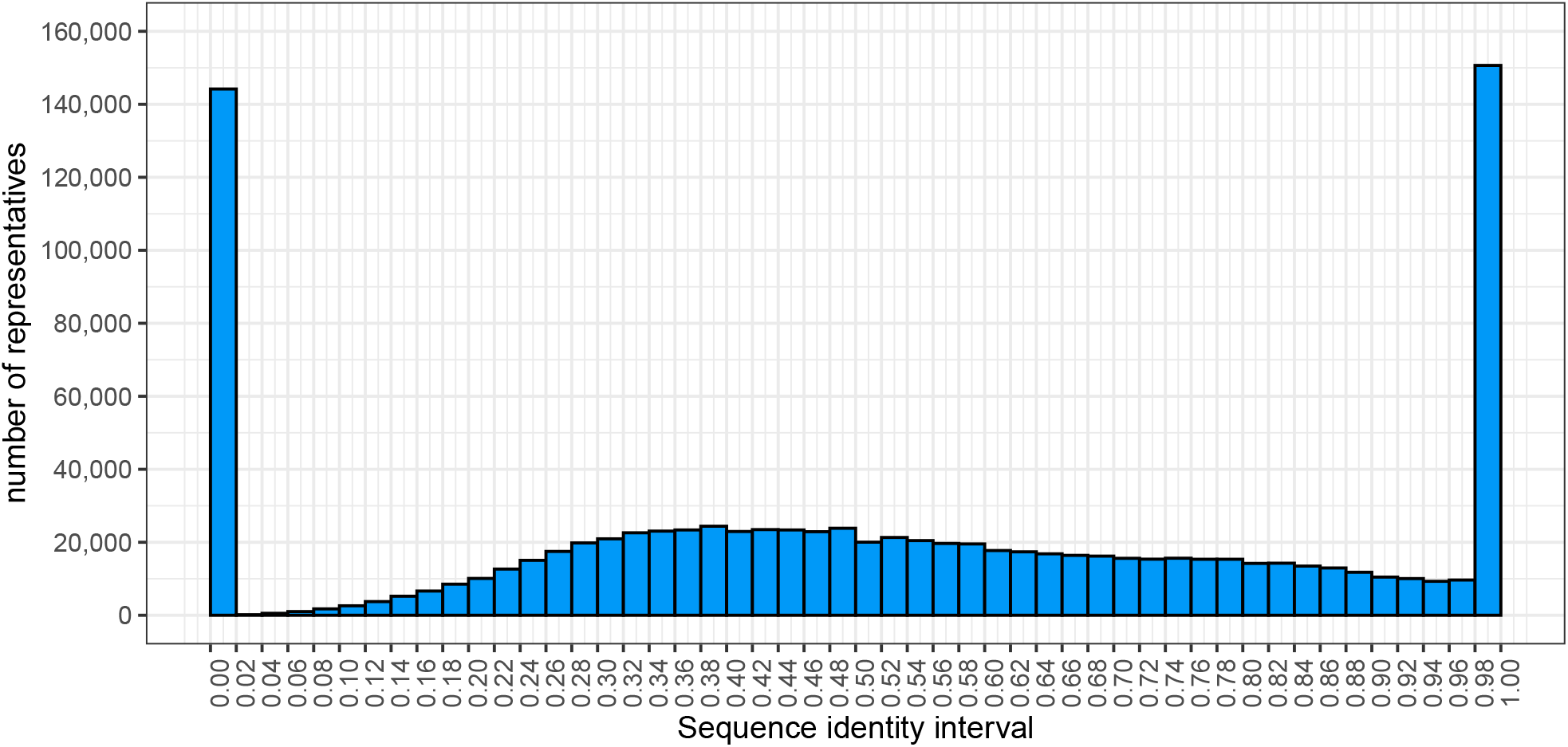
DIAMOND protein search of DIAMOND DeepClust generated representative sequences against public databases. Shown is the sequence identity distribution when aligning a randomly drawn sample of one million representative sequences from clusters of size ≥ 5 from our experimental study against the combined BFD+MGnify databases.

To quantify the relatedness of our clusters to existing protein databases, we sampled one million representative sequences from the clusters containing five or more sequences and aligned them against the combined MGnify^21^ and BFD^6^ databases using DIAMOND in ultra-sensitive mode. We selected the best hit for each query and show the quantity of identities in the alignment divided by the length of the query as a histogram (Supplementary fig. 12) (queries with no hit are assigned zero identities). These results hint toward the possibility that the known protein sequence diversity space of microbes is already fairly saturated today. For the same representative sample, we computed ProtT5^16^ embeddings (using the ProtT5-XL-UniRef50 model) for all sequences of length below 1024 and used mean-pooling of these embeddings as the basis for computing a UMAP^31^ projection (Fig. 2). We labeled sequences according to whether (a) they could be annotated over ≥ 60% of their range against the combined SCOP^19^+ECOD^20^+CATH^17^ databases using DIAMOND in ultra-sensitive mode, (b) they could be annotated over ≥ 60% of their range against the Pfam-A database^18^ using HMMER^32^ at an e-value threshold of 0.001, (c) an alignment against UniProt was found using DIAMOND ultra-sensitive that satisfied an e-value threshold of 0.001, a query coverage threshold of 90% and a sequence identity threshold of 30%, corresponding to the clustering criterions of our clustering and the BFD, (d) an alignment against BFD+MGnify was found satisfying the same criteria, (e) none of the above. If multiple conditions were true, we chose the label according to the order in the previous sentence.

### Data retrieval of the protein biosphere

A total of 22,788,215,153 publically accessible protein sequences were retrieved from JGI IMG^33^, SRC and MERC^34^, MGnify^21^, Metaclust^15^, NCBI NR^35^, AGNOSTOS^36^, MetaEuk^37^, SMAGs^38^, TOPAZ^39^, GPD^40^, NovelFams^41^and MGV^12^ databases during March-April 2022 (additional details are provided in supplementary table 1).

### Data pre-processing and filtering

We first sorted each downloaded FASTA file individually in memory by the length of the sequence in descending order, followed by a global disk-based merge sort on all files using GNU sort, resulting in a combined file of 22,788,215,153 protein sequences. Next, we computed hashes for all sequences which resulted in a deduplicated set of 19,387,935,704 unique sequences.

### Clustering

We clustered the combined input file using a cascaded clustering approach in four rounds at increasing sensitivity employing the DIAMOND modes --faster, --fast, default and --sensitive, also using the option to linearize the comparison in the first round as described in the cascaded clustering section. The clustering criterion was 90% coverage of the cluster member sequence and 30% approximate sequence identity, corresponding to the parameters used to generate the Alphafold2 BFD^6^. To limit the use of resources and create checkpoints that could be reverted to in case of an error, we conducted the clustering rounds as an incremental procedure as follows. First, we split the input sequence file into chunks that we process in sequential steps. Each chunk was first aligned against the current working set of representatives that resulted from the previous steps. Sequences that align against a representative were assigned to its respective cluster. Next, we subjected the remaining sequences that failed to map against an existing representative to all-vs-all alignment at the current round’s sensitivity level and determined new representatives using the greedy vertex cover algorithm, which were then added to the working set. The clustering computation of the unclustered input sequences ran for 6.38 days on a single high-memory 72-core node for the first round, 36.8 hours on 16 nodes for the second round, 20.1 hours on 16 nodes for the third round, and 9.63 days on up to 27 nodes for the fourth round. In total, the computation consumed ∼255,000 CPU hours. The resulting output of DIAMOND DeepClust generated 1,697,446,279 clusters, where 68% of clusters denote singletons and 20% of clusters had three or more members.

## Code availability

DIAMOND DeepClust is available as Open Source Software under the GPL3 license from https://github.com/bbuchfink/diamond.

## Data availability

We will make the Experimental Study dataset freely available upon journal publication. All source datasets that were used are publicly available.

