## Supplementary Information for "Sensitive clustering of protein sequences at tree-of-life scale using DIAMOND DeepClust"

### Main benchmark

#### Environment

All tools were compiled natively from source on the host system using GCC 9.4.0.

#### DIAMOND

We used DIAMOND v2.1.1 available at https://github.com/bbuchfink/diamond[8]. We used the options -DKEEP_TARGET_ID=ON -DHIT_KEEP_TARGET_ID=ON -DEXTRA=ON for compilation, which enable some experimental optimizations not active by default. We implemented the option --mmseqs-compat to improve comparability of raw alignment output with MMSeqs2 by making DIAMOND imitate the alignment score and e-value computation procedure of MMSeqs2. We designed the following commands for the clustering runs:

```
# 90% identity (linear stage only)
diamond deepclust -d nr.dmnd -o out.tsv -M 1000G --member-cover 80 --
cluster-steps faster_lin --approx-id 90 --kmer-ranking --mmseqs-compat
--masking 0 -e 0.00001 --comp-based-stats 0 --swipe-task-size 10000000
--gapopen 10
# 90% identity
diamond deepclust -d nr.dmnd -o out.tsv -M 1000G --member-cover 80 --
approx-id 90 --kmer-ranking --mmseqs-compat --masking 0 -e 0.00001 --
comp-based-stats 0 --swipe-task-size 10000000  --gapopen 10
# 50% identity
diamond deepclust -d nr.dmnd -o out.tsv -M 1000G --member-cover 80 --
approx-id 50 --soft-masking 0 --mmseqs-compat --no-block-size-limit --
masking 0 -e 0.00001 --comp-based-stats 0 --swipe-task-size 10000000
--gapopen 10
# deep clustering
diamond deepclust -d nr.dmnd -o out.tsv -M 1000G --member-cover 80 --
mmseqs-compat --no-block-size-limit --masking 0 -e 0.00001 --comp-
based-stats 0 --swipe-task-size 10000000  --gapopen 10 --anchored-
swipe
# deep clustering (very-sensitive)
diamond deepclust -d nr.dmnd -o out.tsv -M 1000G --member-cover 80 --
cluster-steps faster_lin fast default more-sensitive very-sensitive --
mmseqs-compat --no-block-size-limit --masking 0 -e 0.00001 --comp-
based-stats 0 --swipe-task-size 10000000 --gapopen 10 --anchored-swipe
# deep clustering (ultra-sensitive)
```

```
diamond deepclust -d sensitive_reps -o out.tsv --member-cover 80 -e
0.00001 --cluster-steps ultra-sensitive --masking 0 --soft-masking 0 -
-mmseqs-compat -M 1000G --no-block-size-limit --ext full --swipe-task-
size 10000000 --comp-based-stats 0 --gapopen 10 -g 4000 --no-reextend
--dbsize 169167076291
```

We note that the default 11/1 gap penalties used by MMSeqs2 correspond to 10/1 penalties in the semantic used by BLAST and DIAMOND. For the ultra-sensitive run, we used the representative set of the sensitive run as input and added the runtimes of the two runs. To run the recluster workflow, the following command was run with the output file of the respective clustering run used as input. We added the runtime of the two runs for the purpose of fair reporting.

```
# deep clustering
diamond recluster -d nr.dmnd --clusters out.tsv -o reclust.tsv -M
1000G --member-cover 80 --mmseqs-compat --no-block-size-limit -e
0.00001 --cluster-steps faster_lin fast default more-sensitive --
masking 0 --soft-masking 0 --anchored-swipe --swipe-task-size 10000000
--gapopen 10 --comp-based-stats 0 --more-sensitive --approx-id 0
```

To run the reassignment computation, we used the clustering output of the very-sensitive run and split the set of non-representative sequences into 166 separate files. The following command was run on a cluster of 72-core servers for each of the files with the set of representative sequences used as the database. We added the runtimes of the individual runs multiplied by 72/64 to obtain the runtime on a single 64-core server and also added the time of the original clustering run for the purpose of fair reporting. Based on the output we reassigned every sequence with a hit to the corresponding representative.

```
diamond blastp -q query.faa -d reps.dmnd -o out.tsv -f 6 qseqid sseqid
evalue qcovhsp -k1 --very-sensitive --query-cover 80 -b 1.01 -e
0.00001 --masking 0 -c1 --dbsize 169167076291 --mmseqs-compat --
anchored-swipe --comp-based-stats 0 --swipe-task-size 10000000 --
gapopen 10
```

#### MMseqs2

We used MMseqs2 release 11 available at https://github.com/soedinglab/MMseqs2[23]. Initially, we intended to use the more recent release 13, but noticed serious issues in the context of clustering sensitivity error rates (refer to MMseqs2 GitHub issue #664 for details). For this reason, we reverted to release 11 which did not exhibit this issue. We also noticed that the cascaded clustering workflow did not correctly pass down the --comp-bias-corr option to its subworkflows and that it did not correctly compute e-values with respect to the size of the original input database. We fixed these issues prior to running our benchmarks. We designed the commands for the clustering runs:

```
# 90% identity (Linclust)
mmseqs linclust nr out . --cov-mode 1 -c 0.8 --min-seq-id 0.9 --comp-
bias-corr 0 -e 0.00001 --cluster-mode 2 --alignment-mode 2
# 90% identity
mmseqs cluster nr out . --cov-mode 1 -c 0.8 --min-seq-id 0.9 --comp-
bias-corr 0 -e 0.00001 --cluster-mode 2 --alignment-mode 2
# 50% identity
mmseqs cluster nr out . --cov-mode 1 -c 0.8 --min-seq-id 0.5 --comp-
bias-corr 0 -e 0.00001 --cluster-mode 2 --max-seqs 20/100 --mask 0 --
alignment-mode 2
# deep clustering s=6.0
mmseqs cluster nr out . --cov-mode 1 -c 0.8 -s 6.0 --comp-bias-corr 0
-e 0.00001 --cluster-mode 2 --max-seqs 20/300/1000 --mask 0
# deep clustering s=6.0 (with cluster-reassign)
mmseqs cluster nr out . --cov-mode 1 -c 0.8 -s 6.0 --comp-bias-corr 0
-e 0.00001 --cluster-mode 2 --max-seqs 300 --cluster-reassign 1 --mask
0
# deep clustering s=7.5
mmseqs cluster nr out . --cov-mode 1 -c 0.8 -s 7.5 --comp-bias-corr 0
-e 0.00001 --cluster-mode 2 --max-seqs 1000 --mask 0
```

We attempted a run using -s 6.0 --max-seqs 2000 and a run using -s 6.0 --max-seqs 1000 --cluster-reassign 1 which failed due to running out of temporary disk space (with 2 TB of space available on the machine).
We computed the assignment error correction based on the deep clustering run with sensitivity s=7.5. Since MMseqs2 does not have a dedicated workflow for this purpose, we performed this computation running the search workflow of all non-representative sequences against all representative sequences using this command:

```
mmseqs search query reps out . -s 7.5  --cov-mode 2 -c 0.8 --max-seqs
1000 --comp-bias-corr 0 --mask 0 -e 0.00001
```

We split the query file into 331 chunks and ran the computation separately for each file on a cluster of 72-core nodes. We added the runtimes multiplied by 72/64 to obtain the time for a single 64-core server. Based on the output we reassigned every sequence to the top hit.

#### CD-HIT

We used CD-HIT v4.8.1 available from https://github.com/weizhongli/cdhit[13]. The command for the clustering run:

```
cd-hit -i nr.faa -o out -c 0.9 -n 5 -M 1000000 -d 0 -T 64
```

#### Evaluation

To compute the ground truth for evaluating the error metrics, for each run we aligned a sample of 3,000 cluster members and 3,000 cluster representative sequences respectively against the database of all representatives of that run using the following command:

```
diamond blastp -q QUERY_FILE -d REP_DB -o OUTPUT_FILE -k0 --swipe -
b100 -e 0.1 --masking 0 --dbsize 169167076291 --mmseqs-compat --
gapopen 10 --comp-based-stats 0 -f 6 qseqid sseqid pident length
mismatch gapopen qstart qend sstart send evalue bitscore qcovhsp
scovhsp approx_pident
```

The option --swipe computes a full Smith Waterman alignment against all database sequences and therefore guarantees perfect sensitivity. Here the database size corresponds to the size of the original input database of the clustering runs.
While both Diamond and MMseqs2 use the approximate sequence identity instead of the actual sequence identity in their clustering criterion, both tools also apply heuristics to cluster sequences based on counting identities on diagonals without gapped extension[15]. To account for this fact and not have this behavior cause errors in the evaluation, we count a sequence as correctly clustered if either the approximate identity or the actual identity of the pairwise local alignment is above the threshold value of the respective run. We do not assume an error if only the actual identity is above the threshold for unclustered pairs. We use the same formula as MMseqs2 to compute the approximate identity in order to ensure comparability: let $l_1$ and $l_2$ be the lengths of the aligned ranges in the two sequences and s be the BLOSUM62 raw alignment score, then the approximate identity is defined as $\min(s/\max(l_1, l_2)*0.1656 + 0.1141), 1)$.

We implemented a separate logic to evaluate CD-HIT as this tool does not use a clustering criterion compatible with the logic used by Diamond and MMseqs2. The ground truth was computed using this command:

```
diamond blastp -q QUERY_FILE -d REP_DB -o OUTPUT_FILE -k0 -c1 -e 1 --
masking 0 --dbsize 155806124097 -b6 --motif-masking 0 --ext full --
comp-based-stats 0 -f 6 qseqid sseqid nident qlen score
```

Analogous to CD-HIT, we used the number of identities in the local alignment divided by the length of the cluster member sequence as a distance measure and checked whether this number is above the threshold of 0.9 to accept a sequence as correctly clustered. We define the closest representative sequence as the representative with the highest alignment score that satisfies the clustering criterion.

### Experimental study

The first round of clustering was performed by incrementally processing 109 chunks of the input file. The command for mapping a chunk to the existing representatives:

```
diamond blastp -q REP_DB -d CHUNK_FILE -c1 -k0 --faster -b300 --lin-
stage1 -f 6 qseqid sseqid corrected_bitscore qstart qend sstart send -
-subject-cover 90 --masking 0 --ext banded-slow --soft-masking tantan
--unaligned-targets UNAL_OUT --approx-id 30 --ignore-warnings
```

For technical reasons, we use the representative database as a query file since this choice significantly accelerates the extension computations in cases where there are sufficiently more targets per query than query sequences per target. Here the --lin-stage1 option manually triggers the Linclust-like logic of only comparing against the longest query sequence for groups of identical seeds. We use the --unaligned-targets option to retrieve all targets that did not align against any query, which are then used as input for the following all-vs-all alignment:

```
diamond blastp -q UNALIGNED_SEQS -d UNALIGNED_SEQS -c1 -k0 --faster -
b200 --lin-stage1 -f 6 qseqid sseqid qcovhsp scovhsp
corrected_bitscore --query-or-subject-cover 90 --masking 0 --ext
banded-slow -o SELF_OUT --approx-id 30 --soft-masking tantan --ignore-
warnings
```

We used `diamond greedy-vertex-cover` with the option `--member-cover 90` to compute the clustering based on the alignment output.

We conducted the second round of clustering based on the input file of 4,204,049,109 clusters from the first round, partitioned into 4 chunks, using these commands for mapping and self-alignment:

```
diamond blastp -q CHUNK_FILE -d REP_DB -o OUT --multiprocessing --
parallel-tmpdir TMP_DIR -c1 -b4 --fast --query-cover 90 --approx-id 30
-k1 -f 6 qseqid sseqid corrected_bitscore qstart qend sstart send --
masking 0 --soft-masking tantan --ext banded-slow
```

```
diamond blastp -q UNALIGNED_SEQS -d UNALIGNED_SEQS -o OUT --
multiprocessing --parallel-tmpdir TMP_DIR -c1 -b4 --fast --query-cover
90 --approx-id 30 -k1000 -f 6 qseqid sseqid corrected_bitscore --
masking 0 --soft-masking tantan --ext banded-slow
```

We used `diamond greedy-vertex-cover` with the option `--edge-format triplet` to compute the clustering based on the alignment output. We conducted the third round of clustering based on the input file of 2,223,989,666 clusters from the second round, using this command for self-alignment of the whole dataset:

```
diamond blastp -q INPUT -d INPUT -o OUT --multiprocessing --parallel-
tmpdir TMP_DIR -c1 -b4 --query-cover 90 --approx-id 30 -k1000 -f 6
```

```
qseqid sseqid corrected_bitscore qstart qend sstart send --masking 0 -
-soft-masking tantan --ext banded-slow
```

We used `diamond greedy-vertex-cover` with the option `--edge-format triplet` to compute the clustering based on the alignment output. We conducted the fourth round of clustering based on the input file of 1,906,267,323 clusters from the third run, using this command for self-alignment of the whole dataset:

```
diamond blastp -q INPUT -d INPUT -o OUT --multiprocessing --parallel-
tmpdir TMP_DIR --mp-self -c1 -b1 -k0 --more-sensitive --query-or-
subject-cover 90 --approx-id 30 -f 6 qnum snum qcovhsp scovhsp
corrected_bitscore qstart qend sstart send approx_pident evalue --
masking 0 --soft-masking tantan --ext banded-slow --freq-masking
```

We used `diamond greedy-vertex-cover` with the option `--member-cover 90` to compute the clustering based on the alignment output.

Supplementary table 1 - Public protein sequence datasets that were used for our Experimental Study

| Database name | Number of sequences | Download date/release | URL |
|---|---|---|---|
| IMG Environmental Aquatic Metagenomes | 6,232,188,951 | March 2022 | https://img.jgi.doe.gov/ |
| IMG Environmental Non-Aquatic Metagenomes | 6,851,118,226 | March 2022 | https://img.jgi.doe.gov/ |
| IMG Host-Associated Metagenomes | 1,905,657,909 | March 2022 | https://img.jgi.doe.gov/ |
| IMG Engineered Metagenomes | 801,972,930 | March 2022 | https://img.jgi.doe.gov/ |
| SRC | 2,022,891,389 | March 2022 | http://wwwuser.gwdg.de/~compbiol/plass/current_release/SRC.fasta.gz |
| MGnify | 1,977,479,951 | 2022_05 | https://ebi-metagenomics.github.io/blog/2019/04/10/Protein-database/ |
| metaclust | 1,757,323,526 | March 2022 | https://metaclust.mmseqs.org/2018_06/ |
| NCBI NR | 465,406,186 | April 2022 | https://ftp.ncbi.nlm.nih.gov/blast/db/FASTA/ |
| AGNOSTOS | 427,306,945 | Ver 5 | https://figshare.com/ndownloader/articles/13264769/versions/5 |
| MERC | 292,137,902 | March 2022 | http://wwwuser.gwdg.de/~compbiol/plass/current_release/MERC.fasta.gz |
| MetaEuk | 12,111,301 | 2019_11 | http://wwwuser.gwdg.de/~compbiol/metaeuk/2019_11/ |
| SMAGs | 10,207,435 | v1 | https://www.genoscope.cns.fr/tara/ |
| TOPAZ | 8,405,914 | v1 | https://osf.io/gm564/ |
| GPD | 7,581,807 | April 2022 | ftp.ebi.ac.uk/pub/databases/metagenomics/genome_sets/gut_phage_database |
| NovelFams | 4,587,583 | March 2022 | https://novelfams.cgmlab.org/ |

| | | | |
|---|---|---|---|
| MGV | 11,837,198 | v1.0 | https://portal.nersc.gov/MGV/ |
| **Total** | **22,788,215,153** | | |
